# Whole Brain Polarity Regime Dynamics are Significantly Disrupted in Schizophrenia and Correlate Strongly with Network Connectivity Measures

**DOI:** 10.1101/543751

**Authors:** Robyn L. Miller, Godfrey Pearlson, Vince D. Calhoun

## Abstract

From a large clinical blood oxygen level dependent (BOLD) functional magnetic resonance imaging (fMRI) study, we report several interrelated findings involving transient supra-network brainwide states characterized by a saturation phenomenon we are referring to as “polarization.” These are whole-brain states in which the voxelwise-normalized BOLD (vnBOLD) activation of a large proportion of voxels is simultaneously either very high or very low. The presence of such states during a resting-state fMRI (rs-fMRI) scan is significantly anti-correlated with diagnosed schizophrenia, significantly anti-correlated with connectivity between subcortical networks and auditory, visual and sensorimotor networks and also significantly anti-correlated with contemporaneous occupancy of transient functional network connectivity states featuring broad disconnectivity or strong inhibitory connections between the default mode and other networks. Conversely, the presence of highly polarized vn-BOLD states is significantly correlated with connectivity strength between auditory, visual and sensorimotor networks and with contemporaneous occupancy of transient whole-brain patterns of strongly modularized network connectivity and diffuse hyperconnectivity. Despite their consistency with well-documented effects of schizophrenia on static and time-varying functional network connectivity, the observed relationships between polarization and network connectivity are with very few exceptions unmediated by schizophrenia diagnosis. We also find that the spatial distribution of voxels most likely to contribute to the highly polarized states (polarity participation maps (PPMs)) differs with a high degree of statistical significance between schizophrenia patients and healthy controls. Finally, we report evidence suggesting the process by which the most polarized states are achieved, i.e. the ways that strongly polarized voxel regions extend, merge and recede also differs significantly between patient and control populations. Many differences observed between patients and controls are echoed within the patient population itself in the effect patterns of positive symptomology (e.g. hallucinations, delusions, grandiosity and other positive symptoms of schizophrenia). Our findings highlight a particular whole-brain spatiotemporal BOLD activation phenomenon that differs markedly between healthy subjects and schizophrenia patients, one that also strongly informs time-resolved network connectivity patterns that are associated with this serious clinical disorder.

## I INTRODUCTION

While there is growing attention in human blood-oxygenation dependent (BOLD) functional magnetic resonance imaging (fMRI) studies focused on time-varying connectivity between functional networks [1–13], much less attention is given to the global spatial dynamics of activated brain space induced by and facilitating time-varying network relationships [14, 15]. These ambient whole-brain spatiotemporal phenomena can exhibit strong relationships with disease [14] and provide a wider-angle lens through which to view the results of network-level analyses. We present here one example of a dynamically varying spatial phenomenon that correlates very strongly with schizophrenia diagnosis, exhibits structured and robust effects on observed functional network connectivity (FNC), correlates significantly with occupancy of certain dynamic functional network connectivity (dFNC) state and also exposes very different patterns of spatiotemporal dynamics and organization in schizophrenia patients (SZs) and healthy controls (HCs). The basis of this analysis is an early-stage transformation of the pre-processed BOLD fMRI volume into voxelwise normalized form (vn-BOLD /vn-fMRI). Although every voxel spends exactly a third of its time in the upper (resp. lower, resp. middle) third of its own *intrinsic activation profile*, the proportion of gray-matter voxels simultaneously in the upper (resp. lower) third of their respective activation profiles during given TRs is highly variable. In healthy subjects we find the brain coalesces regularly into states where a high proportion of voxels are simultaneously in the highest (resp. lowest) third of their activation profiles, i.e. states in which the brain is highly *polarized*.

These highly polarized brain states occur significantly more often in healthy controls than in schizophrenia patients (see Figure 1). The more balanced state characterized by an approximately equal proportion of voxels in the highest, lowest and middle thirds of their intrinsic activation profiles (IAPs) occurs much more frequently in schizophrenia patients. These polarized regimes are also very strongly correlated with short-timescale whole-brain functional network connectivity states (some positive, some negative correlations in which the effect direction differs between patients and controls for the highly modularized connectivity state exhibiting negative DMN-to-other connections). The phenomenon of widespread polarization can be parsed more finely in spatial terms, and doing so also reveals distinctions between patients and controls. Examples include the determination of each voxel’s “*polarity participation rate*” which is its probability of being in the top (resp. bottom) third of its own activation profile at times when this condition holds for a large proportion of voxels, i.e. when the whole brain is in a polarized state. We find that the spatial distribution of voxelwise polarity participation differs significantly between patients and controls; in controls there are large clearly delineated regions characterized by very high polarity participation rates, while in patients the larger and better defined regions are those that represent very low rates of polarity participation. Another way to probe the time-varying spatial structure of polarized voxels is to recode (*polarity code*) voxel timeseries as {−1,0,1} according to whether the voxel is in the bottom, middle or top third of its intrinsic activation profile at each TR. The polarity-coded brain maps can then be clustered into a set of brainwide voxel *co-polarization patterns*, providing a spatially explicit parameterization of polarity space in which subject trajectories can be studied. We find highly significant differences between patients and controls in occupancy rates of the resulting clusters.

**Figure 1.**
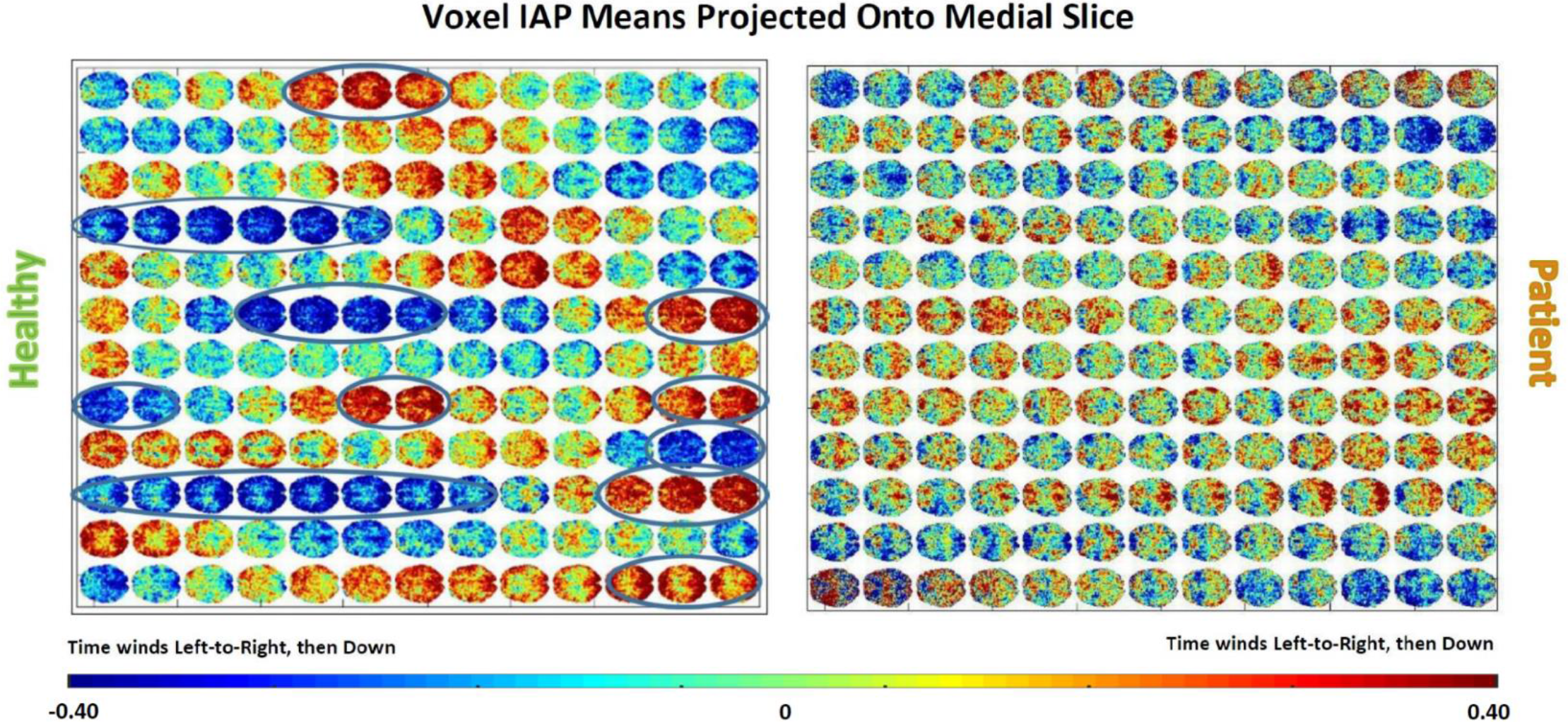
Key observation: healthy brains converge toward and achieve states of pervasive polarization. Both panels show mean normalized voxel timeseries values along the z-dimension displayed on a medial slice advancing through time from left to right; (Left) a 42 yr old healthy male control exhibits intervals in which apparently very large proportions of gray matter voxels are simultaneously toward the top or the bottom of their intrinsic activation profiles (IAPs); (Right) a 40 yr. old male schizophrenia patient exhibits some weak progression toward accumulating large proportions of similarly activated voxels, but the process never quite achieves the highly *polarized* states evident in the matched control displayed on the left.

## II MATERIALS AND METHODS

### A. Participants, Preprocessing, Network Identification and Dynamic Functional Connectivity

We summarize below some basic information about data collection, preprocessing, network identification and computation of dynamic functional connectivity. Details are available in [4]. Resting state functional magnetic resonance imaging data (162 volumes of echo planar imaging BOLD fMRI, TR = 2 sec.) was collected from 163 healthy controls (117 males, 46 females; mean age 36.9) and 151 age and gender matched patients with schizophrenia (114 males, 37 females; mean age 37.8) during eyes closed condition at 7 different sites across United States. After standard preprocessing, the fMRI data from all subjects was decomposed using group independent component analysis (GICA) into 100 network spatial maps (http://icatb.sourceforge.net) of which 47 were identified as functionally meaningful resting state networks (RSNs). Subject specific spatial maps (SMs) and timecourses (TCs) were obtained from the group level spatial maps via spatiotemporal regression. The timecourses were detrended, orthogonalized with respect to estimated subject motion parameters, then despiked and band-pass filtered into [0.05,0.15] Hz using a 5^th^ order Butterworth filter. Dynamic functional connectivity (dFNC) between RSN timecourses was estimated using a sliding window approach. Following protocols from recent studies on dynamic connectivity [4], we used a tapered rectangular window length of 22 TRs (44 seconds), shifted 1TR forward at a time and computed pairwise correlations between RSN time courses within these windows. Static, or traditional functional network connectivity (FNC) simply uses a single untapered window of length 162TR, the length of the scan. The 47 functionally meaningful networks retained from the 100 component GICA all into 5 broad functional domains: subcortical (SC, 5 networks), auditory (AUD, 2 networks); visual (VIS, 11 networks), sensorimotor (SM, 6 networks), cognitive control (CC, 13 networks), default mode network (DMN, 7 networks), cerebella (CB, 2 networks).

Pre-processing of 4D volumes for the polarity analysis was identical to that for dFNC, except for omission of the standard spatial smoothing step. This choice was made to control for cases in which a relatively small number of evenly distributed extremely high (or low) intensity voxels distort polarity measurements for the regions they are distributed within.

### B. Dynamic Polarity States, Polarity Metric and Polarity Participation Maps

We construct several measures intended to capture spatial patterns in the brain formed by the set of voxels that are either simultaneously very highly activated or simultaneously minimally activated. These *co-polarized* voxels do not exclude voxel collections that share correlative relationships, but they are far more general. Included in the set of voxels, for example, that are highly activated during a given TR there would usually be a subset that rose together from an earlier TR, a subset that dropped together from an even higher previous level of activation and a subset that are roughly unchanged from the previous TR. Which is to say that within the set of highly activated voxels at a given TR can be, in a temporally localized sense, subsets consisting of correlated voxels, anti-correlated voxels and uncorrelated voxels.

#### 1) Voxel Activation Profiles and Polarity Coding

The timeseries of each gray matter voxel 𝑣 ∈ 𝒱 is individually *z*-scored to create a normalized (intrinsic) activation profile. Values in each *z*-scored timeseries are then discretized into discrete intrinsic activation level (IALs), i.e. they are recoded as −1s, 0s or +1s according to whether they are in the lower third (*z* < −0.43, ‘Low Polar’), middle third (*z* ∈ (−0.43,0.43), ‘Neutral’) or upper third (*z* > 0.43, ‘High Polar’) of the voxel’s intrinsic activation profile (see Figure 2). The resulting discretized timeseries in 𝒜 = {−1,0, +1} ≡ {*Low*, *Neutral*, *High*} for each voxel will be referred to as the its discretized intrinsic activation level timeseries, *IAL*_𝑣_(*t*).

**Figure 2.**
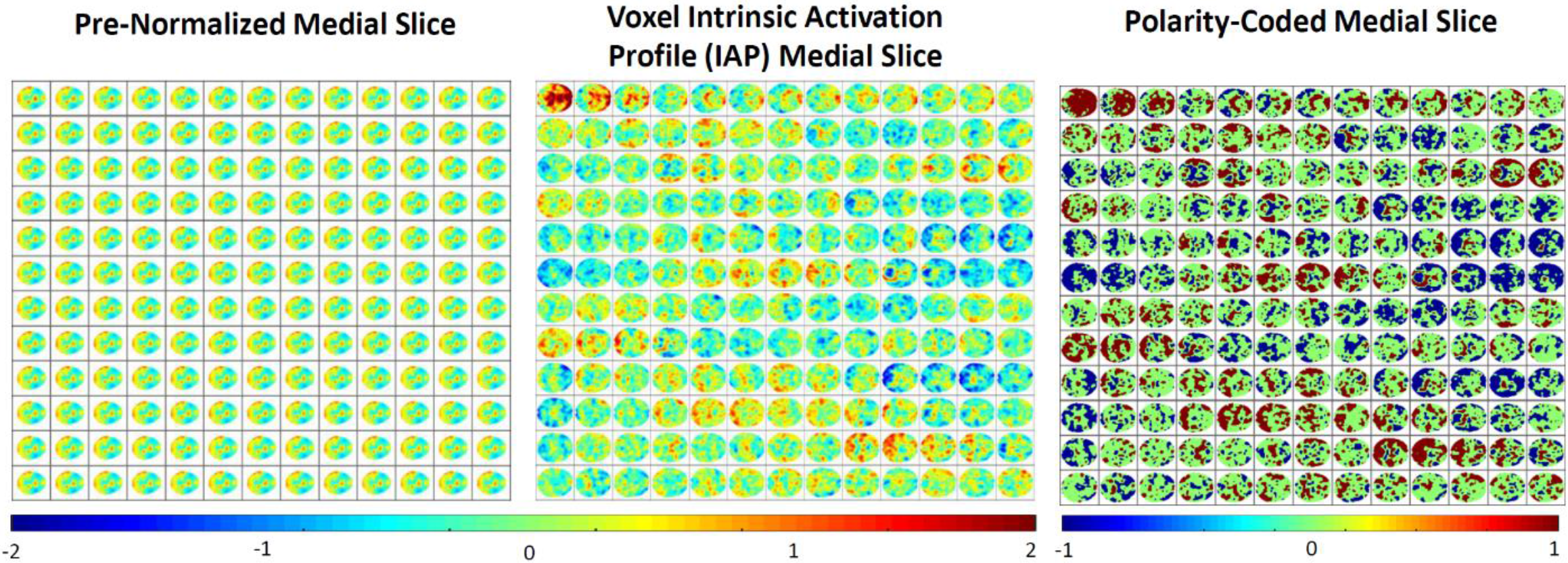
Time advancing first along rows, then along columns in all three panels; (Left) Basic post-processed medial slice data through time for one subject; (Center) Same subject’s medial slice data through time where each voxel’s activation timeseries has been separately z-scored to recast the data in a voxel-intrinsic, normalized form that provides a clearer view of which voxels are near the ceiling or the floor of their own activation profile at each TR]; (Right) Discretized version of the middle panel in which voxel IAPs are recoded in {−1,0,+1} according to whether they are in the bottom third of the IAP (*z* < −0.431), the middle third of the IAP (−0.431 ≤ *z* ≤ 0.431) or top third (*z* > 0.431) as a rough parsing of the brain at each TR into voxels near their own floors and ceilings of activation.

#### 2) Dynamic Polarity Regimes (dPRs)

Every individual voxel intrinsic activation level timeseries assumes, by design, each of the values −1, 0 and 1 exactly 1/3 of the time. This however provides no information about the proportion of voxels simultaneously at the same IAL at any given TR. Switching from the individual voxel standpoint to the whole brain standpoint, we characterize each TR by a length-3 vector 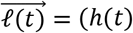, *l*(*t*), *n*(*t*)), *h*(*t*), *l*(*t*), *n*(*t*) ∈ [0,1], *h*(*t*) + *l*(*t*) + *n*(*t*) = 1 of proportions where *h*(*t*) = #{𝑣: *IAL*_𝑣_(*t*) = 1}/|𝒱|, *l*(*t*) = #{𝑣: *IAL*_𝑣_(*t*) = −1}/|𝒱|, *n*(*t*) = #{𝑣: *IAL*_𝑣_(*t*) = 0}/|𝒱| are the proportion of voxels at each intrinsic activation level at TR=*t*. If we do not keep track of the entire history of each voxel, the number of voxels in any of the three intrinsic activation levels observed at a given TR would follow a binomial distribution with *p* = 1/3, *n* = |𝒱|. The expected number for any of the levels would be 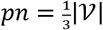 and the expected proportion in each level would simply be 𝔼(*h*), 𝔼(*l*), 𝔼(*n*) = *p* = 1/3. Empirically we see *μ*(*h*) = 0.333, *μ*(*l*) = 0.335, *μ*(*n*) = 0.332. Applying *k*-means clustering (Matlab’s built-in *k*-means function with 3 clusters, 3000 iterates, 300 replicates, Euclidean distance, number of clusters selected by the elbow criterion) to all 50240 of these 3-vectors (one for each of 314 subjects at 160 TRs) suggest the brain dynamically organizes into three very stylized polarity regimes (dPRs) (see Figure 6): a High-Polarized regime whose cluster centroid (0.46,0.24,0.30) presents *h* ≫ 1/3, *l* ≪ 1/3 and *n*~ 1/3; a Low-Polarized regime whose cluster centroid (0.23,0.46,0.31) presents *h* ≪ 1/3, *l* ≫ 1/3 and *n*~ 1/3; and a Non-Polarized or Equidistributed Regime whose cluster centroid (0.33,0.33,0.34) presents *h*~ 1/3, *l*~ 1/3 and *n*~ 1/3 (see Figure 4).

#### 3) Polarity Metric

We also define a real-valued polarity metric. Since 𝔼(*h*), 𝔼(*l*) = 1/3 and the polarized states are characterized by having both *h* > 1/3 *and l* < 1/3, a given subject’s degree of polarization at *TR* = *t* can be summarized with *Π*(*t*) = −(*h*_*z* (*t*)*l*_*z* (*t*)), where *h*_*z*_ *and l*_*z*_ are z-scored versions of the timeseries *h*(*t*) and *l*(*t*) respectively, recentered to have mean 0. *Large* positive values of *h*_*z*_ (resp. *l*_*z*_) represent values of *h*(*t*) (resp. *l*(*t*)) that are much larger than 1/3, while high magnitude negative values of *h*_*z*_ (resp. *l*_*z*_) represent values of *h*(*t*) (resp. *l*(*t*)) that are much smaller than 1/3. The most polarized timepoints are those in which one of *h*_*z*_ *or l*_*z*_ is large and positive, while the other is large and negative. So *Π*(*t*) is rising with the degree of polarization at *TR* = *t*, is near zero when the brain is balanced between *h*(*t*), *l*(*t*) and *n*(*t*) and grows more negative when both *h*(*t*) and *l*(*t*) are simultaneously much larger or much smaller than 1/3 (brain is not polarized but also not well balanced between *h*(*t*), *l*(*t*) and *n*(*t*)) (see Figure 3).

**Figure 3.**
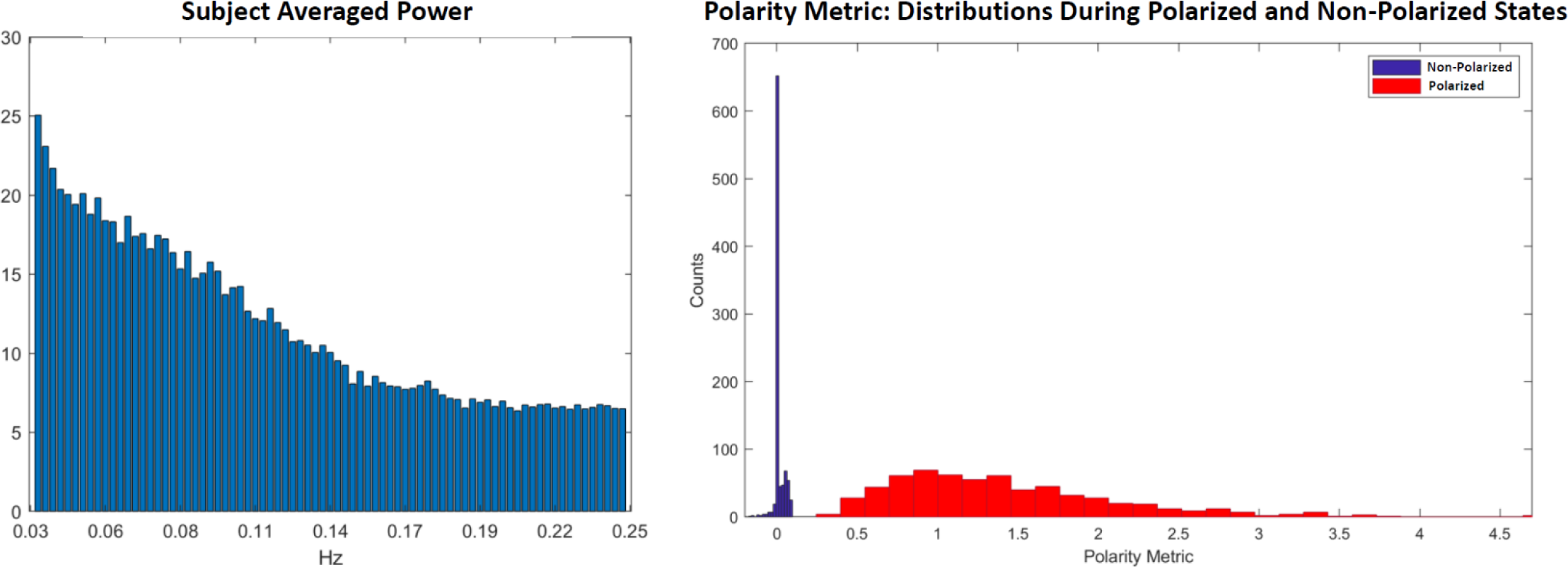
Using scalar valued measure **Π**(***t***) of time-varying whole-brain polarization we see that (Left) the oscillatory pattern in degree of polarity has power decaying in frequency and no discernible peaks that would point to biological rhythms associated with respiration or heartbeat; (Right) The value of **Π**(***t***) clearly separates for timepoints at which the whole brain is in the Non-Polarized dPR state (blue) vs. timepoints at which the whole brain is one of the two polarized (Polarized-High, Polarized-Low) states (red).

### c. Polarity Participation Maps (PPMs)

The dPRs are crude summaries and capture nothing about the locations or identities of voxels that contribute to the computed proportion of voxels in each of the three levels at a given TR, and thus nothing about the overall role of specific voxels in producing the a subject’s polarized regimes. To capture this information we create a polarity participation map (PPM) for each subject in which each voxel is assigned a value in [0,1] according to the proportion of polarized dPR states it contributes to, i.e. the sum of the proportion of that subject’s Polarized-High states during which that voxel’s *IAL*_𝑣_(*t*) = 1 and the proportion of their Polarized-Low states during which the voxel’s *IAL*_𝑣_ = −1 (see Figure 4). Applying *k*-means clustering (Matlab’s built-in *k*-means function with 2 clusters, 3000 iterates, 300 replicates, Euclidean distance, number of clusters selected by the elbow criterion) to all 314 of these 60303-vectors (one for each of 314 subjects) yields clusters whose centroids evidently represent different spatial distributions of highly participating voxel (Figure 9, bottom left).

**Figure 4.**
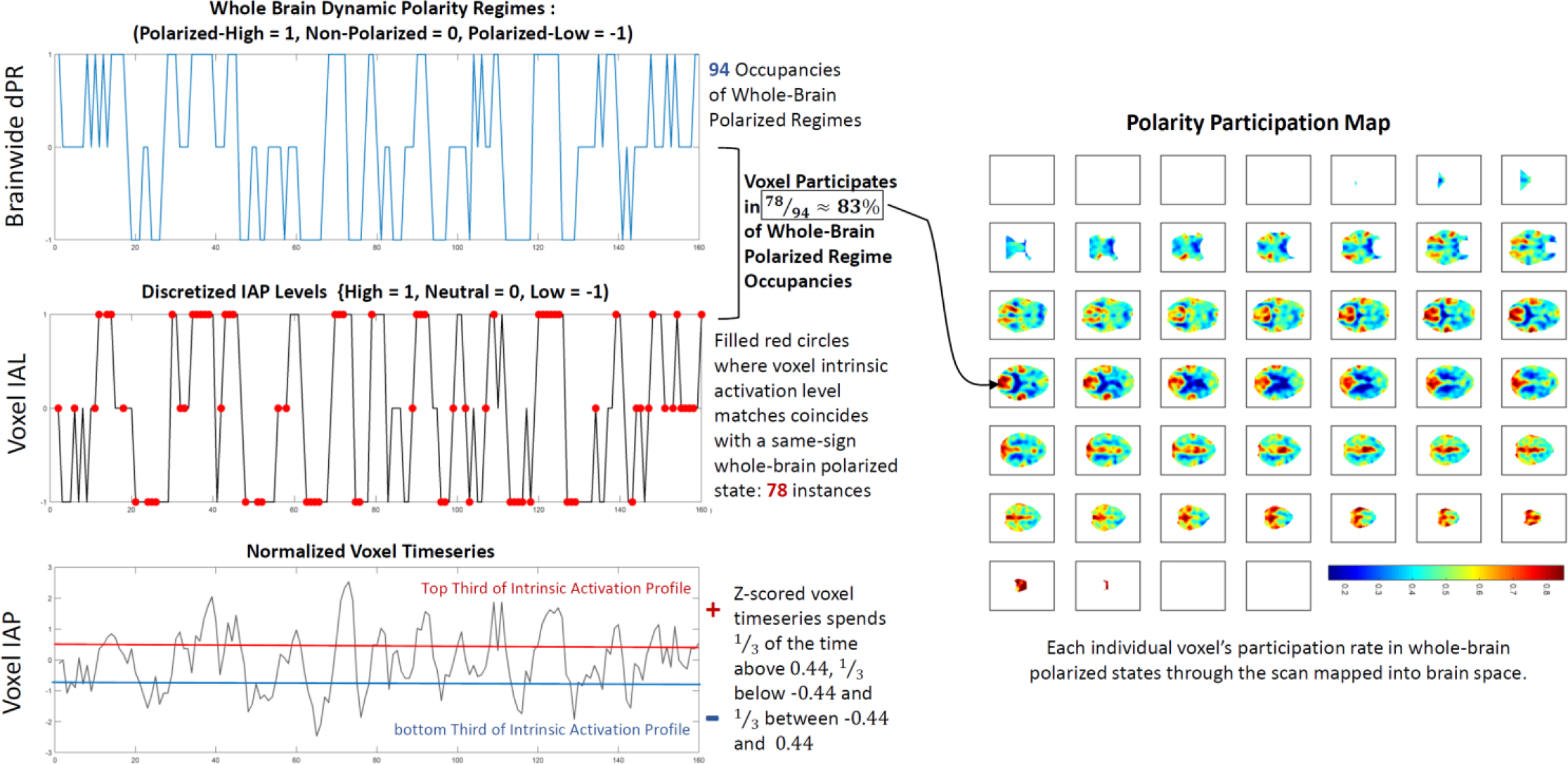
Overview schematic showing the steps from (Bottom Left) individually normalized voxel timeseries (voxel intrinsic activation profiles (IAPs)) to (Middle Left) single voxel discretized intrinsic activation levels (IALs) to (Top Left) whole brain dynamic polarity regimes (dPRs) into which this voxel is recruited at a very high rate (83% of the time that the whole brain is in the Polarized-High or Polarized-Low state, each characterized by mean of of the entire brain’s voxels are simultaneously either in the high or low (+1, −1) intrinsic activation levels, this voxel is one of the participating set of co-polarized voxels. The average voxel only participates in fewer than half (43%) of a subject’s highly polarized states.

#### 1) Network-Level Polarity Participation

To assess the role that different functional networks play in whole-brain polarized states, we threshhold and binarize the 47 group-level network spatial maps and then average the value of each subject’s *z*-scored polarity participation map restricted to the threshold-surviving voxels in each network spatial map (see Figure 10, bottom row). Specifically, each group spatial map is z-scored (since scale in GICA spatial maps is arbitrary) and thresholded at *z* ≥ 1.25 to retain the only the top decile of voxels. The polarity participation rates are also *z-*scored and averaged, for each subject, over each network’s threshold-surviving voxels. The measured network participation for a given subject and a given network can thus be negative, indicating that the average voxel in that network participates in the subject’s whole-brain polarized states at a sub-mean level.

### D. Dynamic Brainwide Co-Polarization Patterns (CoPPs)

The low-dimensional dPRs give a rough sense of how overall counts of co-polarized voxels vary through time and PPMs provide a time-averaged picture of which regions of the brain are most commonly implicated in whole-brain polarized states. Neither approach, however, provides insight into how highly polarized whole-brain states evolve and dissolve, nor what role they may play in longer whole-brain itineraries that involve more structured or intricate spatial patterns of polarity-coded voxels. Toward this end we extract transient whole-brain patterns of co-polarization in a voxel-intrisic analogy to whole-brain co-activation patterns (CAPs) employed in several published studies [16]. Replacing each continuous voxel timeseries 𝑣(*t*) with its discretized intrinsic activation level timeseries *IAL*_𝑣_(*t*) yields a representation of the entire brain in 𝒜 = {−1,0, +1} ≡ {*Low*, *Neutral*, *High*} at each TR. Applying *k*-means clustering (Matlab built-in *k*-means function with 13 clusters, 3000 iterates, 100 replicates, Euclidean distance, number of clusters selected by the elbow criterion) to all 50240 of these |𝒱| = 6030-dimensional observations (one for each of 314 subjects at 160 TRs) spatially elaborates the crude numerical counts/proportions captured by the 3-dimensional dynamic dPRs. The brainwide co-polarization patterns (CoPPs) that emerge as centroids of these clusters indicate that the cruder dPRs are capturing spatially structured voxel-level arrangements whose population-level occupancy rates are relatively uniform (Figure 11, top left). Among the CoPPs associated with TRs that are assigned to one of the polarized dPR states are some that are pervasively polarized, i.e., the proportion of voxels simultaneously at either High or Low levels is much higher than the ~45% or so in the associated dPR cluster centroid (see Figure 11, bottom left).

#### 1) Identification of the most ‘Strongly Polarized’ CoPPs

Although the samples being clustered are 60303-dimensional vectors of −1’s 0’s and +1’s, the elements of the cluster centroids are averages in the continuous interval [−1, 1]. Due to the high dimensionality of the samples (i.e., membership in a given cluster can arise predominantly from good matching of some arbitrary collection of tens of thousands of voxels to other cluster members, while still leaving thousands of degrees of freedom), the determination of what it means for one of these centroids to represent a “pervasively polarized” brain state is necessarily somewhat subjective. We have decided on a criterion based loosely on the asymptotic distributions of sample means for a set of identically distributed uniform discrete random variables under a convenient (but not entirely valid assumption) of independence. The sample is quite large so the asymptotic framing is not entirely unreasonable. We will say that a CoPP represents a strongly polarized state if the probability that the magnitude of the absolute mean |*μ*_*CoPP*_| of its elements would have probability less than *α* = 0.001 under a very conservative adaptation of above-stated model. The sign of *μ*_*CoPP*_ in such a case determines if that CoPP is positively or negatively polarized. Since voxels assume values +1,0 ans −1 with equal probability, i.e. 𝔼(*IAL*_𝑣_(*t*)) = 0, 𝔼((*IAL*_𝑣_(*t*) − 𝔼(*IAL*_𝑣_(*t*))^2^) = 1, we assume by the central limit theorem that the within-cluster voxel means are distributed as 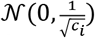 where *c*_*i*_ is the number of observations in the cluster, and thus the global mean *μ*_*i*_ of the whole set of 𝑣 = 60303 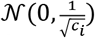 random variables is distributed as 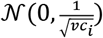. The rescaling factor 𝑣*c*_*i*_ > 10^7^ for all *i*, which makes the standard deviation so close to 0 that absolute values as small as10^−5^ are in the extreme *α* = 0.001 tails of the distribution. To account for the high level of spatiotemporal dependence and produce a distribution whose far tails do not include values with magnitude smaller than 10^−5^, we first reduce *c*_*i*_ by treating all timepoints from a given subject as dependent, so that cluster *i* is treated as containing *n*_*i*_ = 314*r*_*i*_ samples, where *r*_*i*_ is the population-averaged occupancy rate of cluster *i*. We further assume, based on the group spatial ICA (GICA) performed on this data [4] that there are 47 independent units of space (the set of “good” spatially independent networks identified by the GICA). The test distribution for *μ*_*i*_ thus becomes 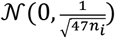, a distribution with much more conservative tails than 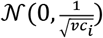. Using the *α* = 0.001 tails of this distribution as a threshold we identify 3 positively (see Figure 5) and 3 negatively polarized CoPPs

**Figure 5.**
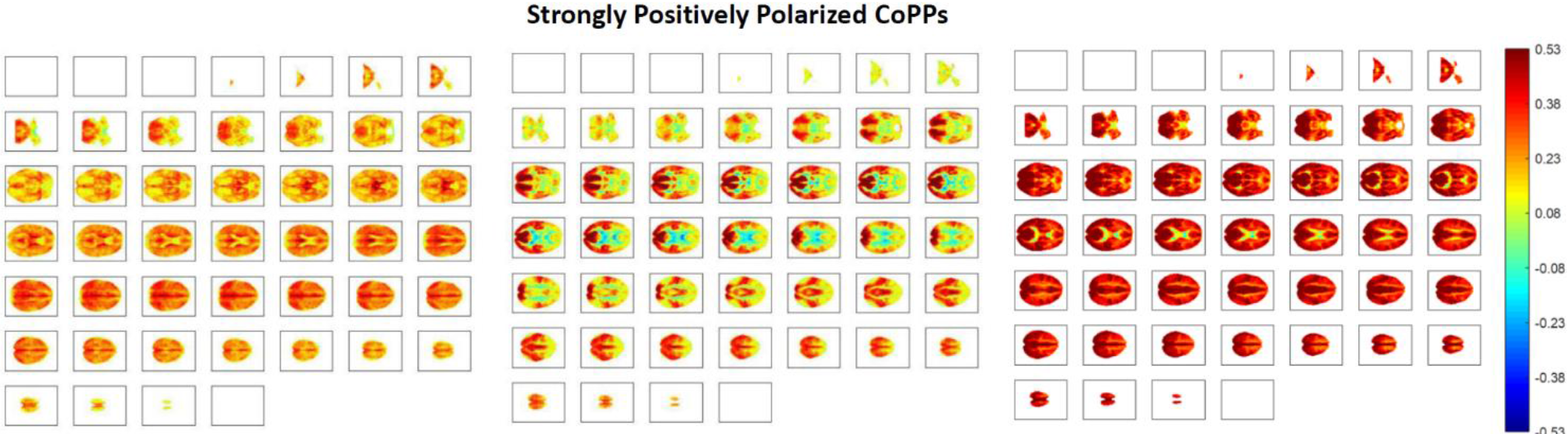
All panels show axial slices from *z* = 1 (top left corner) to *z* = 46 (bottom right corner); (Left) one of the strongly polarized transient time varying co-polarization patterns (CoPPs); (Middle) more structured but still strongly positively polarized co-polarization pattern; (Right) extremely strongly and pervasively positively polarized CoPP.

### E. Longer Itineraries in Polarity Space

We take two approaches to investigating forward-time CoPP itineraries:

#### 1) Itineraries from Group-Averaged Transition Probabilities

We find the forward-time group-level most-probable itinerary from a source CoPP *C*_*i*_ through a set of subsequent *distinct* CoPPs *C*_*j*_, *C*_*k*_, *C*_*l*_, … where *i* ≠ *j* ≠ *k* ≠ *l* ≠ … by using transition probabilities computed at the group level: from *C*_*i*_ the most probable forward-time transition out of *C*_*i*_ at the group-averaged level is to *C*_*j*_, and from *C*_*j*_ the most probable transition is to *C*_*k*_ etc. This process which can be extended until one of the elements of the path repeats, at which point the remainder of the forward-time itinerary is determined by the segment that contains the initial instance and the successor instance of the repeated element. Note that these are itineraries with respect to CoPP transitions from one state to *a new distinct* state, not accounting for the probability of remaining in the presently occupied CoPP. For purposes of interpretation it is important to keep in mind both that the reported itineraries are computed from group-level average transition probabilities and that they are expressed in terms of successive distinct states rather than successive timepoints. We are employing this approach to better understand how healthy and patient populations move through the polarity space (parameterized in terms of CoPPs) and specifically to probe the role the most pervasively polarized states may be playing in overall brain function.

#### 2) Sparse Convolutional Coding of CoPP Occupancy Timeseries

Another way to study longer sequences of CoPP occupancies is to perform multichannel 1-dimensional convolutional sparse coding (CSC) of the symbolic timeseries of CoPP occupancies. These are symbolic timeseries because although we are indexing the CoPPs with a set of nonnegative integers, the actual index values have no numerical meaning: they can be permuted, replaced with large negative numbers, replaced with letters or words or any other way of identifying a given CoPP with some tag or label. This is a slightly unconventional type of timeseries to approach with convolutional feature extraction techniques, however it can be handles by treating the 1D symbolic timeseries in a 𝑁-letter alphabet as an 𝑁-channel binary 1D timeseries, which is effectively a multivariate timeseries in which *exactly one* of the channels (or univariate consitutent timeseries) takes value 1 at every timestep, while all others are 0 at that timestep. This encodes the occupancies in channel form and allows the use of convolutional sparse coding algorithms, such as those developed for the the publicly available Python-based sparse coding package Sporco [17]. Based on initial exploratory work, resource/efficiency considerations and concern with ease of interpretation we used Sporco’s convolutional basis pursuit denoising (CBPDN) dictionary learning algorithm to represent the CoPP symbolic timeseries data with a dictionary of 64 mutichannel (in this case 13 channels, one for each CoPP) elements of temporal duration 11TRs. The elements represent replicable translation-invariant features of the mutichannel timeseries data that combine additively at each timepoint to reproduce the local properties of the signal. The elements of the dictionary are also, as an entire set, optimized so that only a small number of the elements are making the largest proportion of nonzero contributions to signal reconstruction, e.g. they are translation-invariant features, built from a nonlinear feature extraction algorithm, that ultimately reproduce the data additively with the largest rate of nontrivial contribution coming from a subset of the features.

### F. Statistical Analyses

All reported schizoprenia or symptom effects from regression analyses were obtained from mutiple regressions in which age gender and mean frame displacement (motion) were included as nuisance variables, i.e. the reported effects are already corrected for age, gender and motion. In the case of symptom effect, the reported results for a specific category of symptom, e.g. positive symptoms of schizophrenia, are also corrected for confounding effects of the remaining symptom categories, e.g. negative and general symtoms.

## III RESULTS

### A. Dynamic Polarity Regimes and Schizophrenia

We find that occupancy of very polarized brain states is strongly correlated with diagnostic status (see Figure 6): SZ diagnosis has a significant negative effect on occupancy of both the Polarized-High (*β* = −0.05, *p* < 1.1 × 10^−13^) and the Polarized-Low (*β* = −0.04, *p* < 2 × 10^−11^) states. Higher scores on a symptom inventory [18] further suppress polarity, with positive symptoms (including psychotic symptoms such as delusions and hallucinations) exhibiting significant suppressive effects on the Polarized-High (*β* = −0.03, *p* < 0.003) and Polarized-Low (*β* = −0.02, *p* < 0.05).

**Figure 6.**
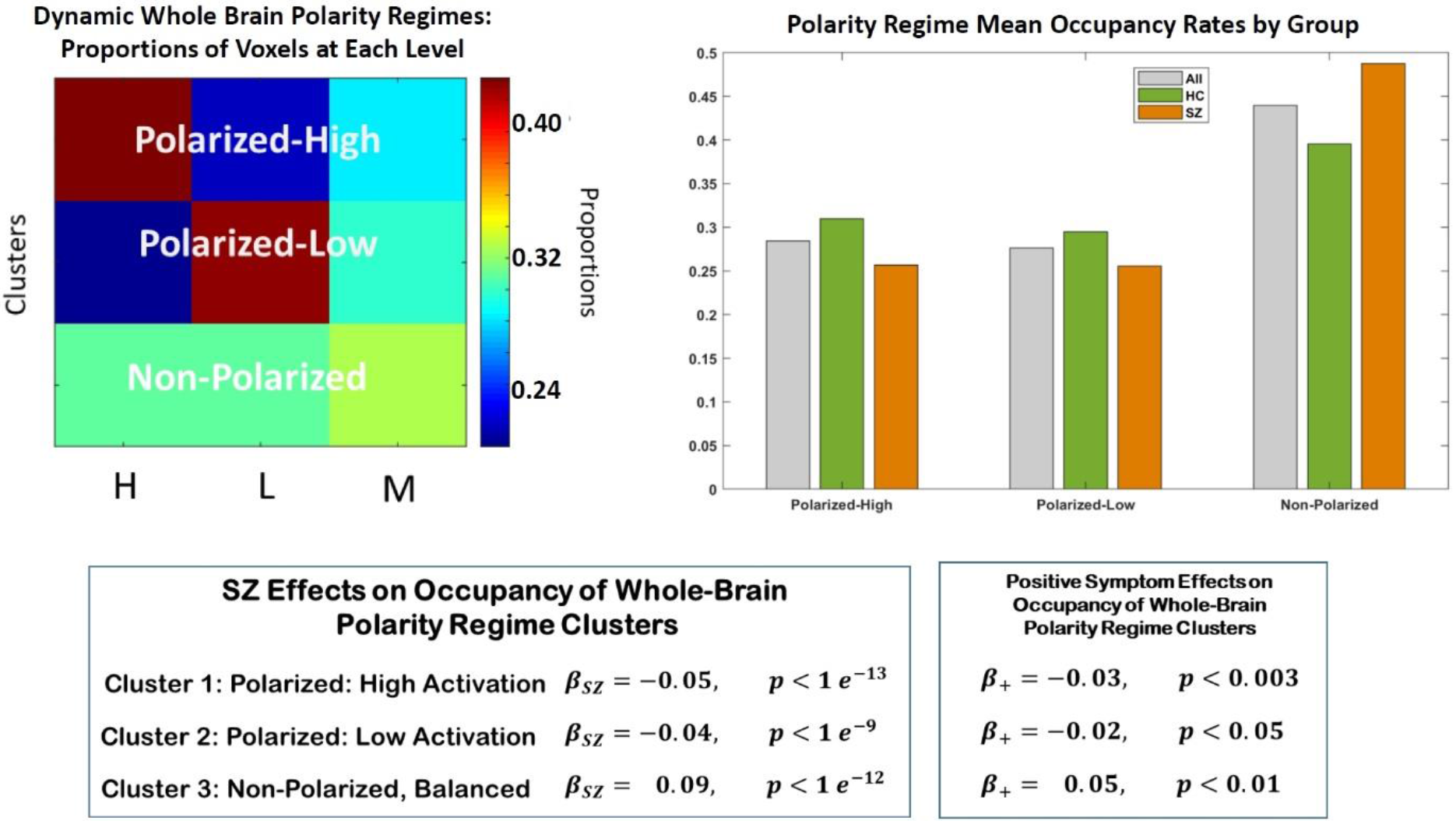
(Top Left) The three whole-brain dynamic polarity regimes: Polarized-High (46% voxels high, 24% voxels low, 30% voxels neutral), Polarized-Low (23% voxels high, 46% voxels low, 31% voxels neutral), Non-Polarized/Balanced (33% voxels high, 33% voxels low, 34% voxels neutral); (Top Right) Group-wise mean occupancy rates of the three dynamic polarity regimes; the non-polarized/balanced regime is the most common for all groups, with evident differences between patients and controls in all regimes; (Bottom Right) In regression model correcting for age, gender and mean frame displacement (motion), the effect of schizophrenia diagnosis on occupancy of the polarized states is negative and highly significant while the effect of schizophrenia diagnosis on occupancy of the balanced state is positive and highly significant; (Bottom Right) Among schizophrenia patients, the effect of positive symptoms (including delusions, hallucination and grandiosity) intensity on polarity regime occupancies echoes the broad effect of schizophrenia itself suggesting that the difference between schizophrenia patients and controls the neighboring may be driven largely by positive symptomology, including the key psychotic symptoms: delusions and hallucinations. Negative schizophrenia symptoms such as bunted affect and general psychological symptoms such as anxiety and depression did not have significant effects on the polarity regime occupancy rates.

### B. Dynamic Polarity Regimes and Functional Network Connectivity

Since the original decomposition into networks is typically based on scale-invariant measures such as correlations (seed-based methods) or dependence (independent component analysis (ICA)), conventional scan-length functional network connectivity (FNC) and also dynamic functional network connectivity (dFNC) computed on sliding windows through the network timecourses both implicitly incorporate information from what we are calling voxel activation profiles. These initial decompositions are followed by evaluations of network connectivity that also tend to rely on correlation, a scale-invariant quantity. Using the five time-resolved connectivity states reported in earlier work from this dataset [19] we probe the relationship between polarity and time-varying connectivity by investigating the occupancy rates of the polarized dPR states within windows characterized by each of the five different dynamic connectivity states (see Figure 7). For each dFNC state, we perform a two-sample *t*-test on the subject-level occupancy rate of polarized dPR states during windows characterized by that dFNC state vs. subject-level occupancy rates of polarized dPR states on all windows. This is assessing group-wise, the degree to which polarized dPR states occupancies during the 22TR windows characterized by any specific dFNC state are distinguished from other 22TR windows in a subject’s scan. The polarized dPR occupancy rates are at the subject-level, computed window-wise for each of the 22TR windows characterized by a given dFNC states for that subject. The number of such windows per subject will vary, so the number of polarized dPR occupancy rates recorded for different subjects for any given dFNC state can be different, and when this count differs between two subject the the number of occupancy rates recorded for those subjects from windows characterized by any other dFNC state will also be different. This somewhat cumbersome procedure allows for testing the role of polarized dPRs in realizing each dFNC state, *correcting for* the tendency of a given subject to be in polarized dPR states more generally. And since the *t*-tests are done at the group level, i.e. they are tests of the entire set of polarized dPR occupancy rates associated with windows characterized by a given dFNC state for all healthy controls (resp. all SZ patients) vs. the entire set of polarized dPR occupancy rates associated with windows characterized by all other dFNC states for all healthy controls (resp. all SZ patients) it is accounting for the different baseline polarized dPR occupancy rates for each group HCs have a higher baseline occupancy rate for the polarized dPR states. We find that for both SZs and HCs, polarized dPR state occupancy is significantly elevated during occupancy of the hyperconnected dFNC state and the highly modularized dFNC state in which the default mode network (DMN) is non-correlated with other networks. The opposite holds for the disconnected dFNC state and the semi-modularized dFNC state in which the DMN is anti-correlated with other networks: polarized dPR state occupancy is significantly depressed during occupancy of these two dFNC states. The highly modularized dFNC state with negative DMN-to-other connectivity is the one for which polarity plays a very different role in the patient and control populations. Polarized state occupancy during occupancy of this dFNC state is significantly elevated in controls, and significantly depressed in patients (see Figure 7 for all effects of polarity on dFNC occupancy, evaluated separately for SZ and HC).

**Figure 7.**
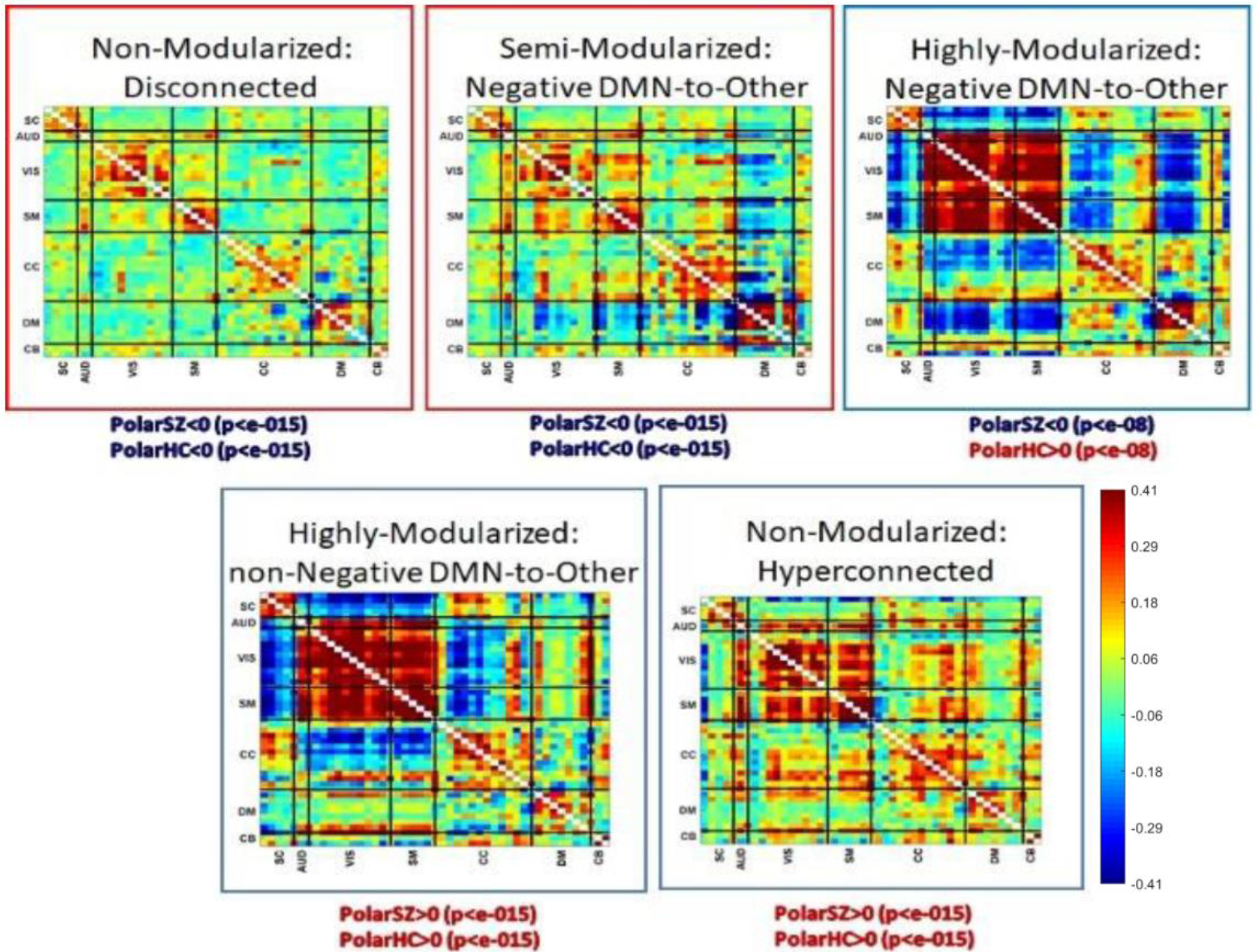
Labeled dFNC states with functional domains indicated along axes: Subcortical (SC), Auditory (AUD), Visual (VIS), Sensorimotor (SM), Cognitive Control (CC), Default Mode (DM), Cerebellar (CB); the dFNC states that are significantly more (resp. less) occupied by SZs are in red (resp. blue) boxes; significant elevation (red) or depression (blue) of brain polarity during windows characterized by each dFNC state are underneath each state.

We also investigated the role of polarized dPR occupancy rates on traditional “static” functional network connectivity. In this case we use the standard full-scan dPR occupancy rates employed in most other sections of this paper, i.e. for this analysis the polarized dPR occupancy rate for a given subject is number of occupancies of either the high or low polarized state for that subject, divided by the total number of TRs) and consider their effect on pairwise network connectivity measures also computed as correlations between network timecourses over the whole length of the scan. Our findings suggest that many reliably observed differences between schizophrenia patients and controls in functional network connectivity may actually be strongly mediated through the role of whole-brain polarization, that the ability of the brain to organize itself into pervasively polarized states (a phenomenon that can have implications for connectivity but correlative relationships can exist within and across bands of intrinsic voxel activation levels) influences the measured correlative relationships that we observe in SZ and HC populations, and also proxies for underlying neurobiological and neurovascular facts that play strong roles in organizing network interactions and convergence into transient highly polarized states. Schizophrenia diagnosis is very significantly negatively correlated with occupancy of polarized dPR states, but in a multiple regression of FNC measures on schizophrenia diagnosis *and* polarized dPR state occupancy rate (along with the usual covariates: age, gender and mean frame displacement), the strong univariate effects of polarized dPR occupancy rate on FNC measures largely survive correction for the role of SZ diagnosis while the strong univariate effects of SZ diagnosis are highly disrupted, leaving only a small set of significant SZ effects after polarized dPR occupancy rates are accounted for (see Figure 8).

**Figure 8.**
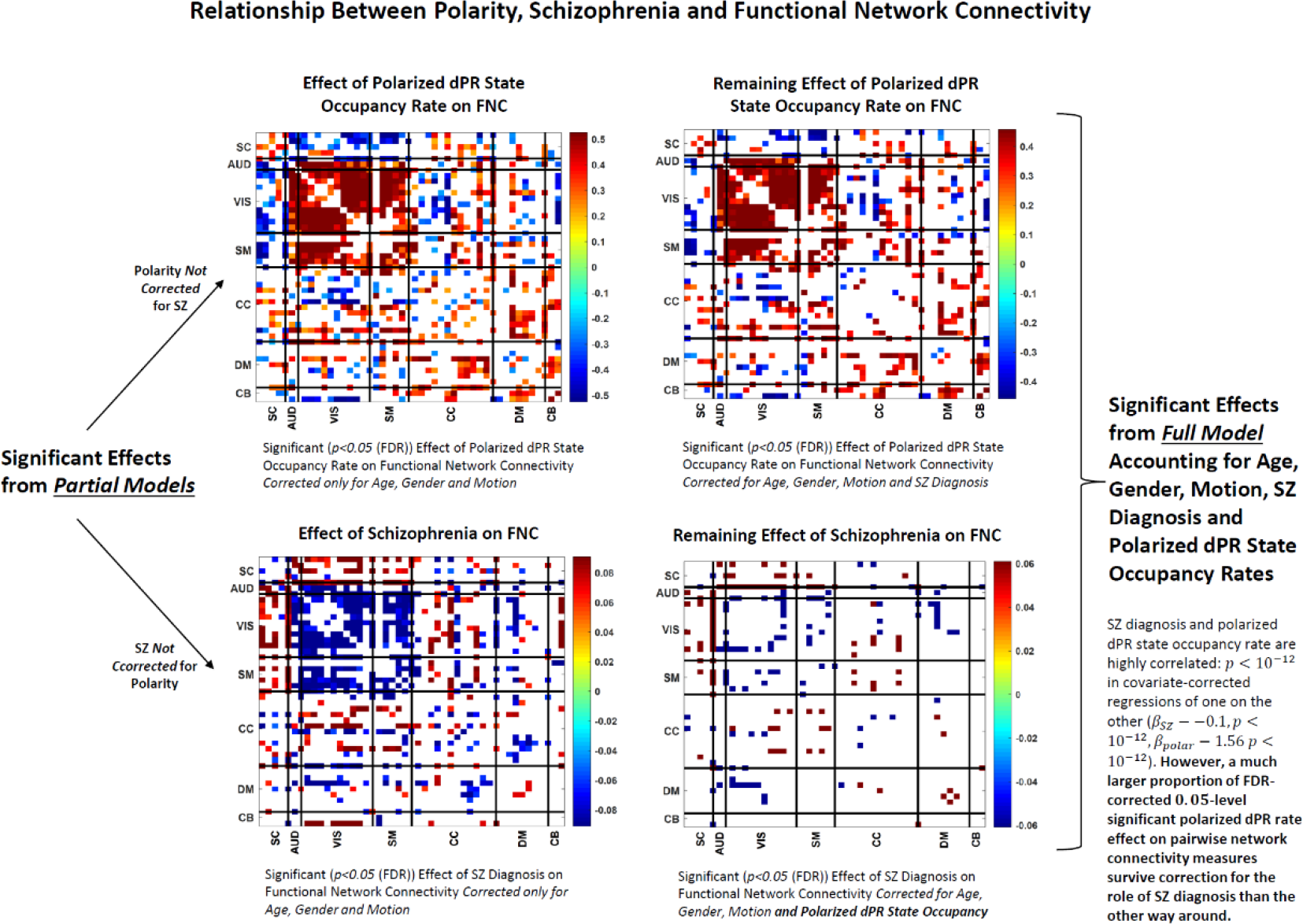
(Top Left) Significant (*p* < 0.05 (FDR)) effects of polarized dPR occupancy on functional network connectivity in multiple regression that accounts for age, gender and motion but not for schizophrenia diagnosis; many strong positive effects within and between auditory (AUD), visual (VIS) and sensorimotor (SM) networks combined with some strong negative effects between subcortical (SC) and cerebellar (CB) networks and the auditory-visual-sensorimotor (AVSM) block; (Top Right) Significant (*p* < 0.05 (FDR)) effects of polarized dPR occupancy on functional network connectivity in multiple regression that accounts for age, gender and motion and also schizophrenia diagnosis; many effects from the partial model to the left survive; broad effect patterns similar after correcting for the role of schizophrenia diagnosis; (Bottom Left) Significant (*p* < 0.05 (FDR)) effects of schizophrenia on functional network connectivity in multiple regression that accounts for age, gender and motion but not for polarized dPR state occupancy rates; many strong negative effects in the AVSM block networks combined with strong positive effects between subcortical and cerebellar networks and the AVSM block (Bottom Right) Significant (*p* < 0.05 (FDR)) effects of schizophrenia on functional network connectivity in multiple regression that accounts for age, gender and motion and also polarized dPR state occupancy; a small proportion of effects from the partial model to the left survive after correcting for the role of dPR state occupancy.

### C. Polarity Participation Maps and Schizophrenia

While schizophrenia patients (especially those with strong positive symptomology are much more likely to pass through periods during which a large proportion of voxels are simultaneously near the ceiling or floor of their own intrinsic activation profiles, the number of ways that proportion of voxels can distribute over space, even subject to local smoothness constraints, is combinatorially overwhelming. For example there are more than 60,303^30,151^ = 6 · 10^4·30,150^ = 6 · 10^121,400^ sets of 30,151 voxels that could be responsible for a given case of the “High-Polarized” state in which ~50% of the voxels are simultaneously at their intrinsic “high” level. Even subject to local smoothness constraints, the participating voxel collections could be drifting spatially between instantiations of the Polarized-High state, yielding relatively uniform, spatially unstructured, time and subject-averaged voxel participation rates. We found however that, on average, there are well-delineated regions in which the most highly participating voxels are concentrated and other regions consisting of voxels unlikely to be part of a whole-brain Polarized-High or Polarized-Low state. Within the broad patterns of voxel participation that hold at the population level, there are also significant differences between schizophrenia patients and controls (Figure 9, top right). The subject-level PPMs robustly fell into two clusters with distinct patterns of voxel participation and highly significant differences in occupancy between patients and controls. The two clusters of PPMs present very different spatial patterns of voxel participation in polarized dPR states. Among other evident differences there are default mode, parietal, cingulate and visual regions that participate in whole-brain polarization at very high rates in Cluster 1 but contribute little to whole-brain polarization in Cluster 2 (see Figure 9, bottom row). The occupancy of both PPM clusters is strongly affected by the diagnostic status of the subject: SZ has a highly significant negative effect (*β* = −0.294, *p* < 2.6 × 10^−08^) on occupancy of Cluster 1 (resp. a highly significant positive effect (*β* = 0.294, *p* < 2.6 × 10^−08^) on occupancy of Cluster 2).

**Figure 9.**
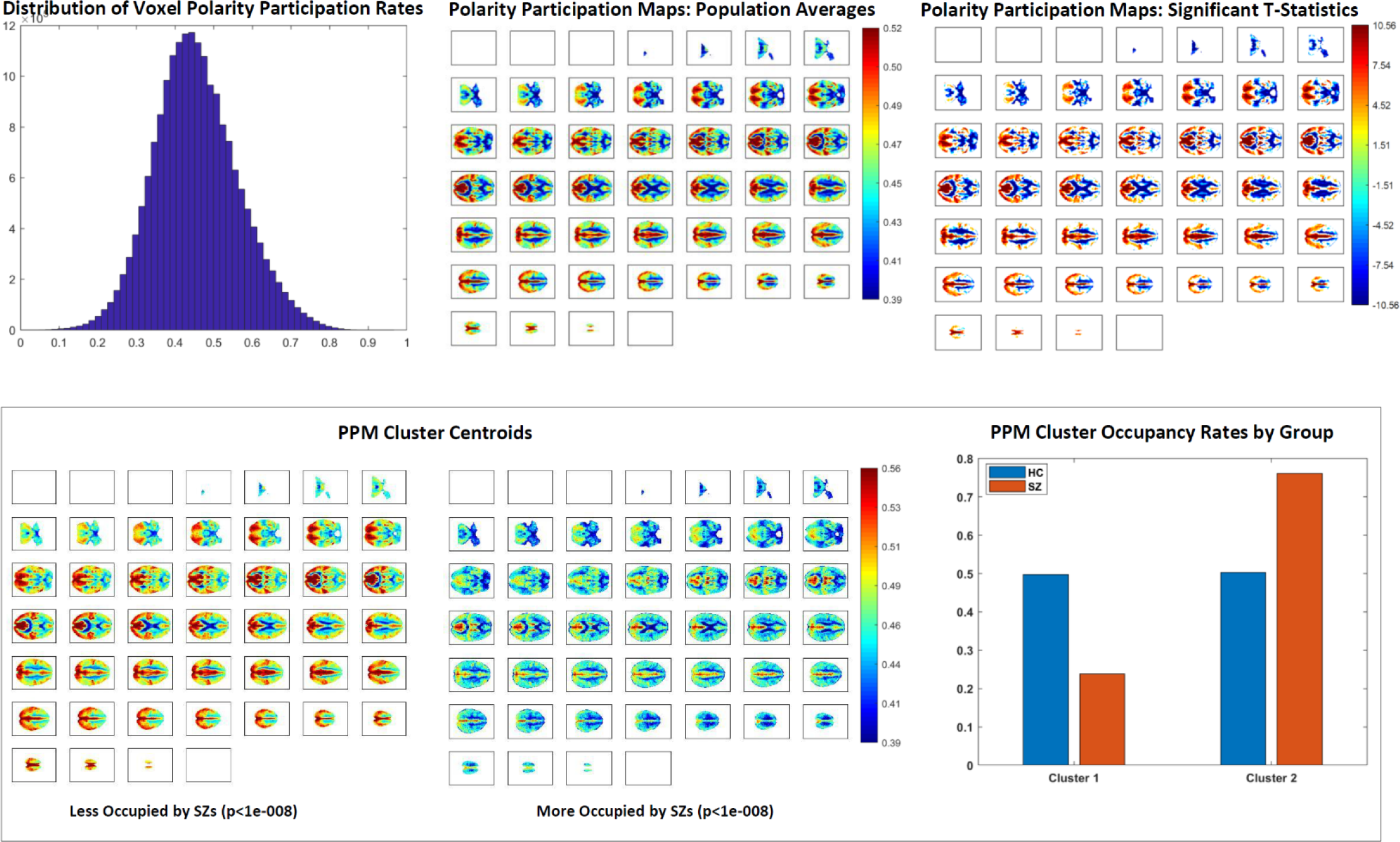
(Top Left) Population-wide voxel polarity participation rates are normally distributed with mean *μ* = 0.46 and standard deviation *σ* = 0.11; (Top Middle) Population average voxelwise polarity participation rates displayed on axial slices from *z* = 1 on upper left through *z* = 46 lower right corner; (Top Right) Significant (*p* < 0.05 (FDR)) voxelwise t-statistics for difference from the mean polarity participation rate; (Bottom Left and Middle) Subject-level polarity participation maps fall robustly into two clusters. Cluster 1 presents regions with above average participation rates, average participation rates and, predominantly ventricular areas with below average participation. The regions with above average participation are much smaller in Cluster 2 than Cluster 1 and the areas with average and below average participation much larger in Cluster 2; (Bottom Right) Healthy subjects split evenly between the two clusters while schizophrenia patients occupy Cluster 2 much more often than Cluster 1.

### D. Polarity Participation Maps, Functional Networks and Schizophrenia

The highest magnitude voxels in GICA functional network spatial maps are, by construction, nearly mutually spatially disjoint. This permits easy assessment of network-level polarity participation by computing the mean polarity participation rate of supra-threshold voxels in each network spatial map. We chose to threshold *z*-scored network spatial maps at *z*̅ = 1.25 (*p*(*z* > *z*̅) < 0.1) and also *z*-scored the PPMs, mapping them from [0,1] into (−∞, ∞). The lowest network-level polarity participation was in subcortical networks (other than the thalamus), the inferior temporal and frontal gyri and the dorsomedial prefrontal cortex (see Figure 10). Significant effects of SZ diagnosis on the network-level polarity participation (*p*<0.05, after false discovery rate (FDR) correction of multiple comparisons) were broadly evident, with positive effects clustering in subcortical cognitive control networks, negative effects in the auditory, visual and sensorimotor networks (see Figure 10). These effects track in sign and significance with SZ effects on inter-network connectivity (see Figure 8 above).

**Figure 10.**
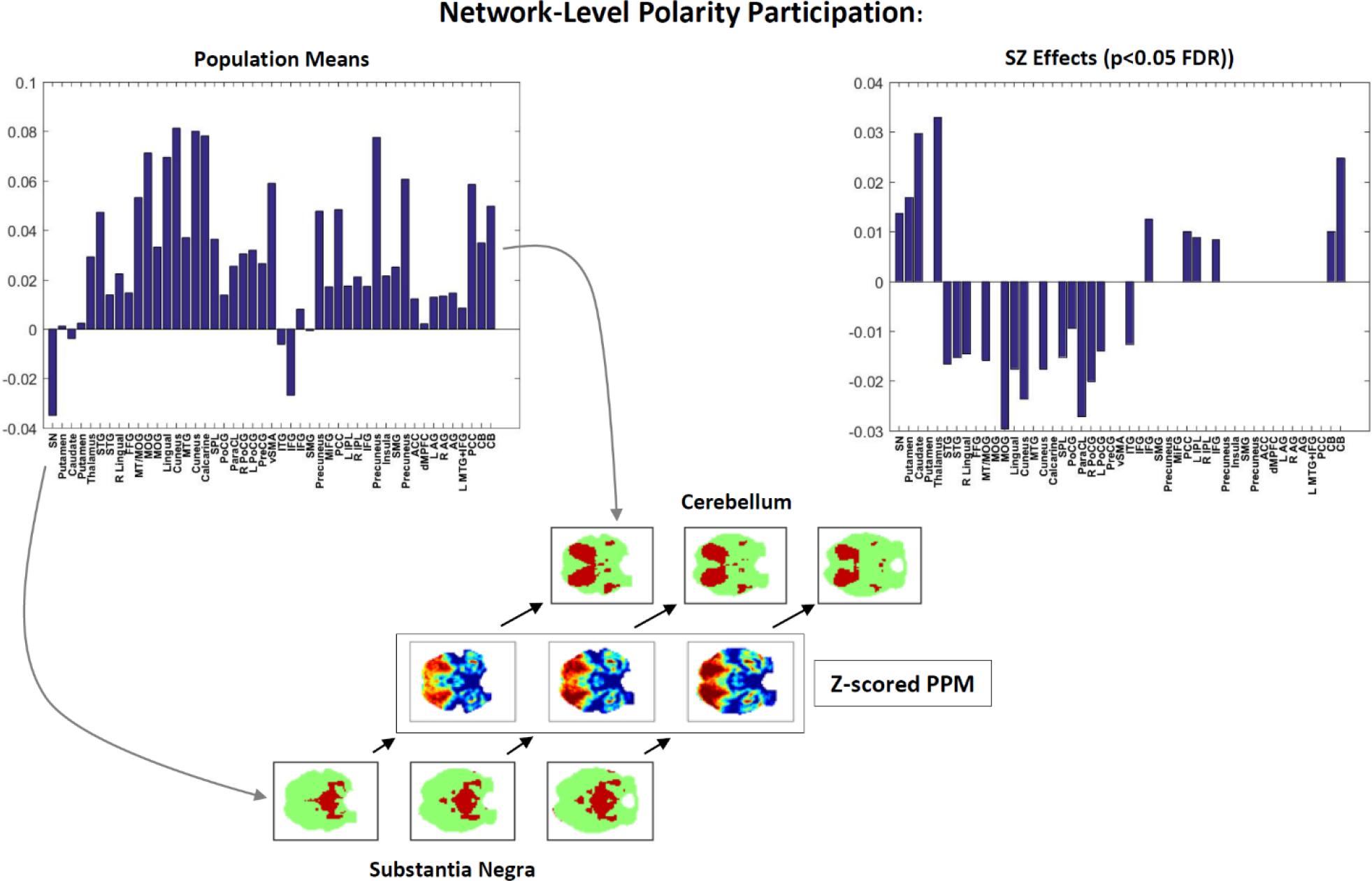
Network level polarity participation; network-level polarity participation population means (top left); significant SZ effects (*p* < 0.05 (FDR)) on network level polarity participation (top right); schematic (bottom) showing three axial slices of the z-scored population averaged PPM (middle) and the same three axial slices of the binarized thresholded group-level cerebellum GICA spatial map (above) and the group-level GICA substantia negra (below) illustrating how PPM projection onto thresholded substantia negra presents negative network-level polarity participation and PPM projection onto thresholded cerebellum presents positive network-level polarity participation.

### E. Dynamic Brainwide Co-Polarization Patterns and Schizophrenia

Brainwide co-polarization patterns, the centroids of clustered 3D polarity-coded volumes (e.g. U_𝑣∈𝒱_ IAL_v_(*t*) for TR=*t*) from all subjects and timepoints, are much more spatially resolved than the dPR *3*-vectors and more temporally resolved than the scan-length summary PPMs. We find that these CoPPs are quite spatially structured, include strong and spatially pervasive polarized states (see Figure 11) and at the population scale, all are occupied at similar rates (see Figure 11). Occupancy of more polarized CoPPs exhibit strong negative SZ effects however.

**Figure 11.**
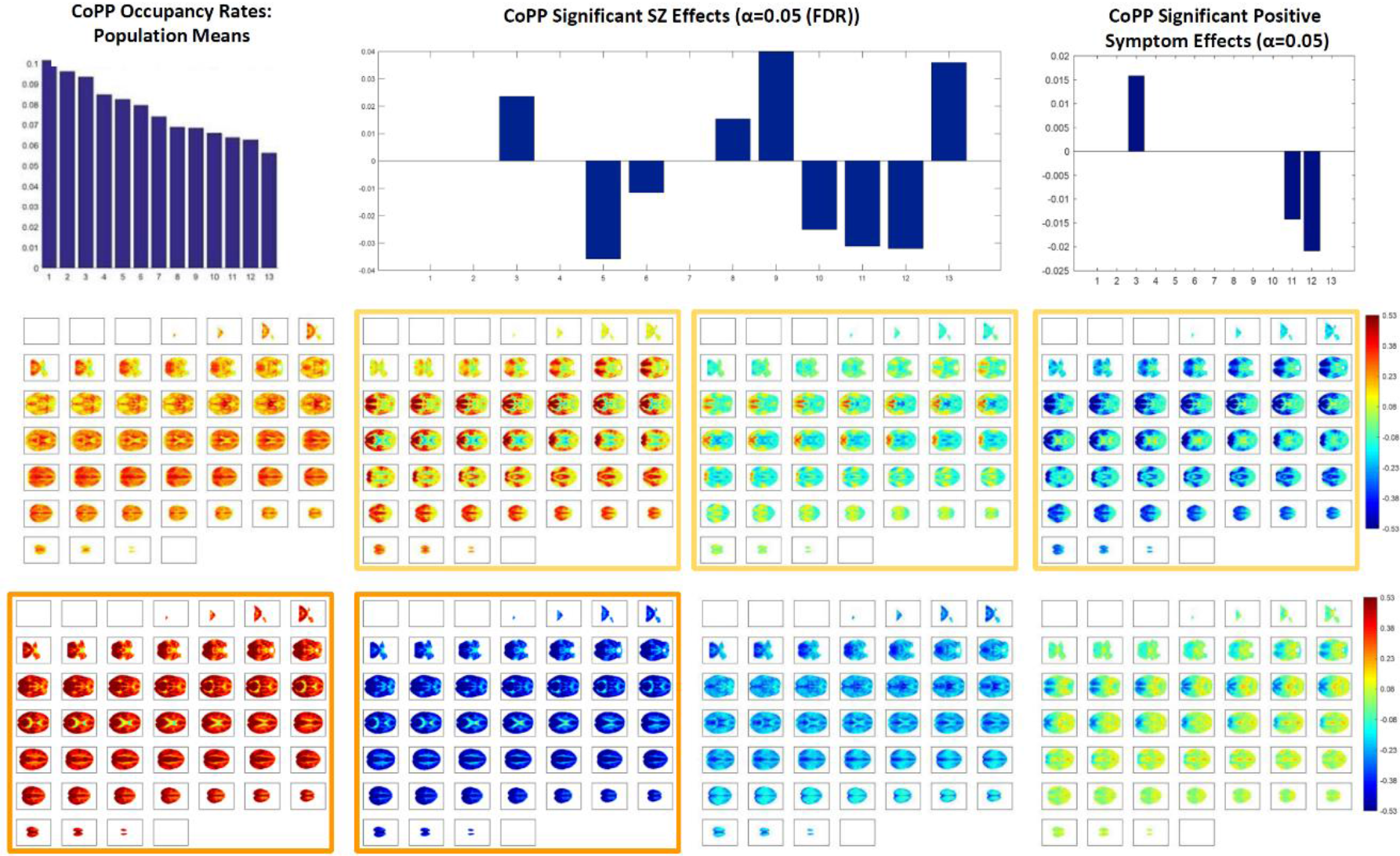
Polarity coded 3D volumes distribute into 13 clusters, all of which exceed 6% occupancy at the population level (top left); among these clusters are two whose centroids are pervasively strongly polarized (bottom row, first and second column); SZ has a significant negative (*p* < 0.05 (FDR)) effect on occupancy of the two most strongly polarized CoPPs (bars at 𝑥 = 11 and 𝑥 = 12 in top-center bar plot); within the patient population there is also a significant negative effect of positive symptomology, including the key psychotic symptoms (delusions and hallucinations) on occupancy of the most strongly and pervasively polarized states (right-most bars in top right bar plot). The positive symptom effect within the patient population is significant at the raw *α* = 0.05 level without surviving the stronger multiple comparison threshold.

#### 1) Longer Itineraries of Co-Polarization Patterns and Schizophrenia

Group-average CoPP transition probabilities reveal that schizophrenia patients move through polarity space, on average, very differently than controls. Considering the longer itineraries in which the most pervasively polarized CoPPs are embedded reveals that their greater probability of realization in healthy controls reflects, in part, a more fluid and diverse sampling of the space of whole brain polarization patterns (Figure 12). Controls exhibit a greater “dynamic range” than patients, a finding consistent with several other recent results [20]. In this setting however, the extra dynamic range puts controls repeatedly in parts of the polarity space that are proximal to more pervasive forms of both positive and negative polarization, increasing the odds of repeatedly entering those states. In healthy controls the most probable exit itinerary from any CoPP has a short transient of at most one intermediate state, followed by entry into the absorbing length-9 periodic orbit joining the most strongly polarized states of both valencies. In patients, the most probable exit itinerary from any CoPP passes through at most four intermediate states before entering one of two absorbing periodic orbits, each an alternating oscillation between either two non-polarized CoPPs or one non-polarized and one mildly polarized CoPP. Some care is required in interpreting these different group-level itineraries: 1. They are expressed in terms of *successive distinct states* rather than successive timepoints; 2. The transition probabilities are group averages that need not be quantitatively or ordinally realized in individual subjects; 3. These are statistical itineraries based on stepwise highest probability transitions and need not be perfectly realized in any fixed set of simulated CoPP occupancy timeseries generated by the full transition probability matrix for either group, i.e. there is non-negligible probability at each step of doing something other than staying in the present state or transitioning to the most probable subsequent distinct state, which makes these itineraries more analogous to trendlines though a pointcloud than to the attractors of a (stochastically perturbed) deterministic dynamical system.

**Figure 12.**
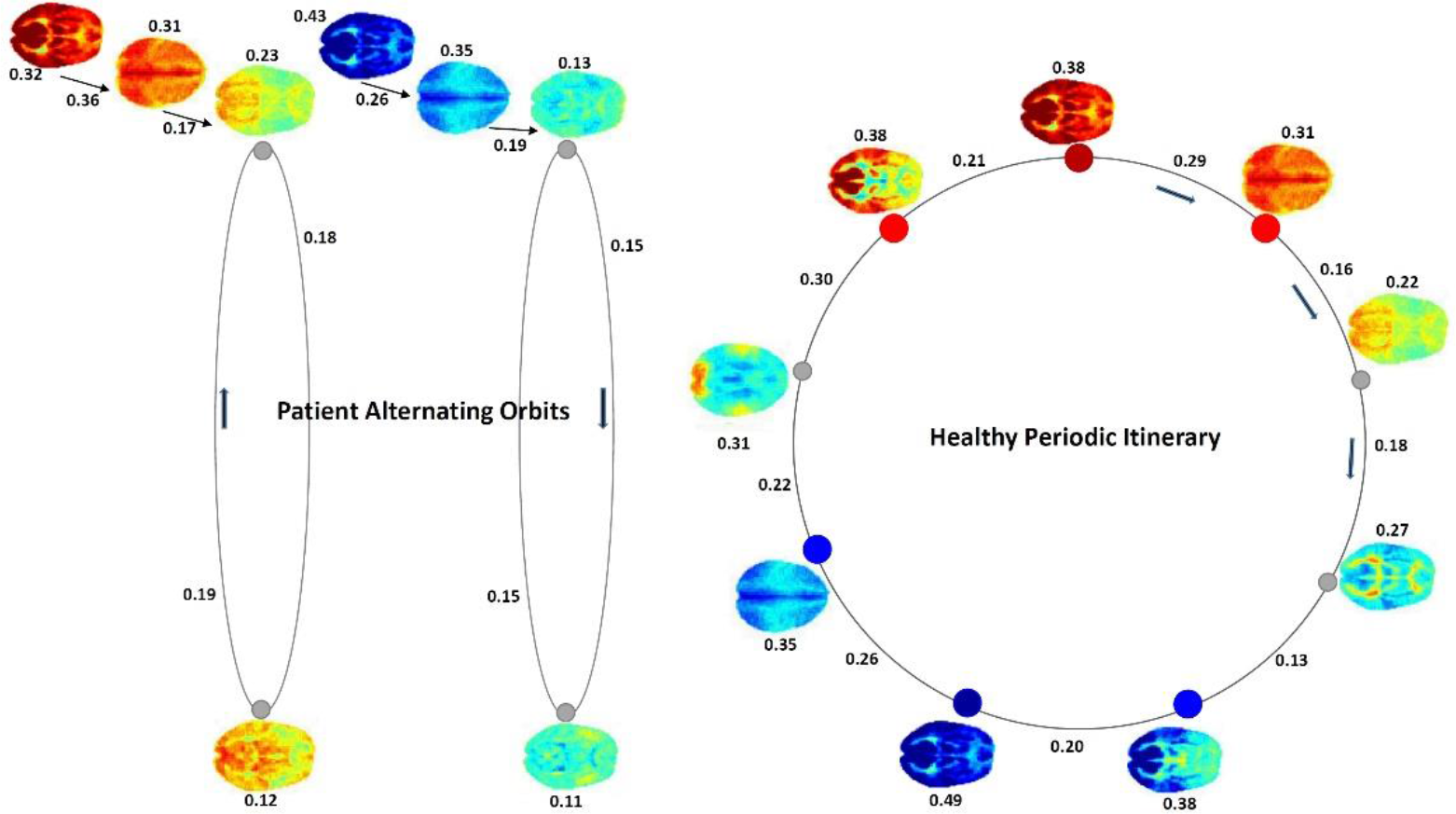
The group level most-probable itineraries out of the maximally pervasively polarized CoPPs differs between controls and patients; Group-specific CoPP-to-self transition probabilities are indicated above or below the CoPP while transition probabilities from one CoPP to the (group-specific) most probable next *distinct* CoPP are indicated along the corresponding edges; highly polarized states are indicated with red (positive) and blue (negative) nodes; (Right) In controls the exit path from either maximally polarized CoPP is the same closed circuit of length 9 which presents a process by which pervasive polarization starts to attenuate, passing through more neutral or heterogeneous states and finally shifting toward pervasive polarization of the opposite valency; After short transients, the most-probable healthy itineraries always enter the displayed length-9 periodic orbit. (Left) In patients, the most probable exit itinerary from either maximally polarized CoPP involves a short transient followed by an alternating oscillation between in one case two neutral non-polarized states, and in the other a negatively polarized state and a non-polarized state. After short transients, the most-probable patient itineraries always enter one of the two displayed alternating oscillations.

**Figure 13.**
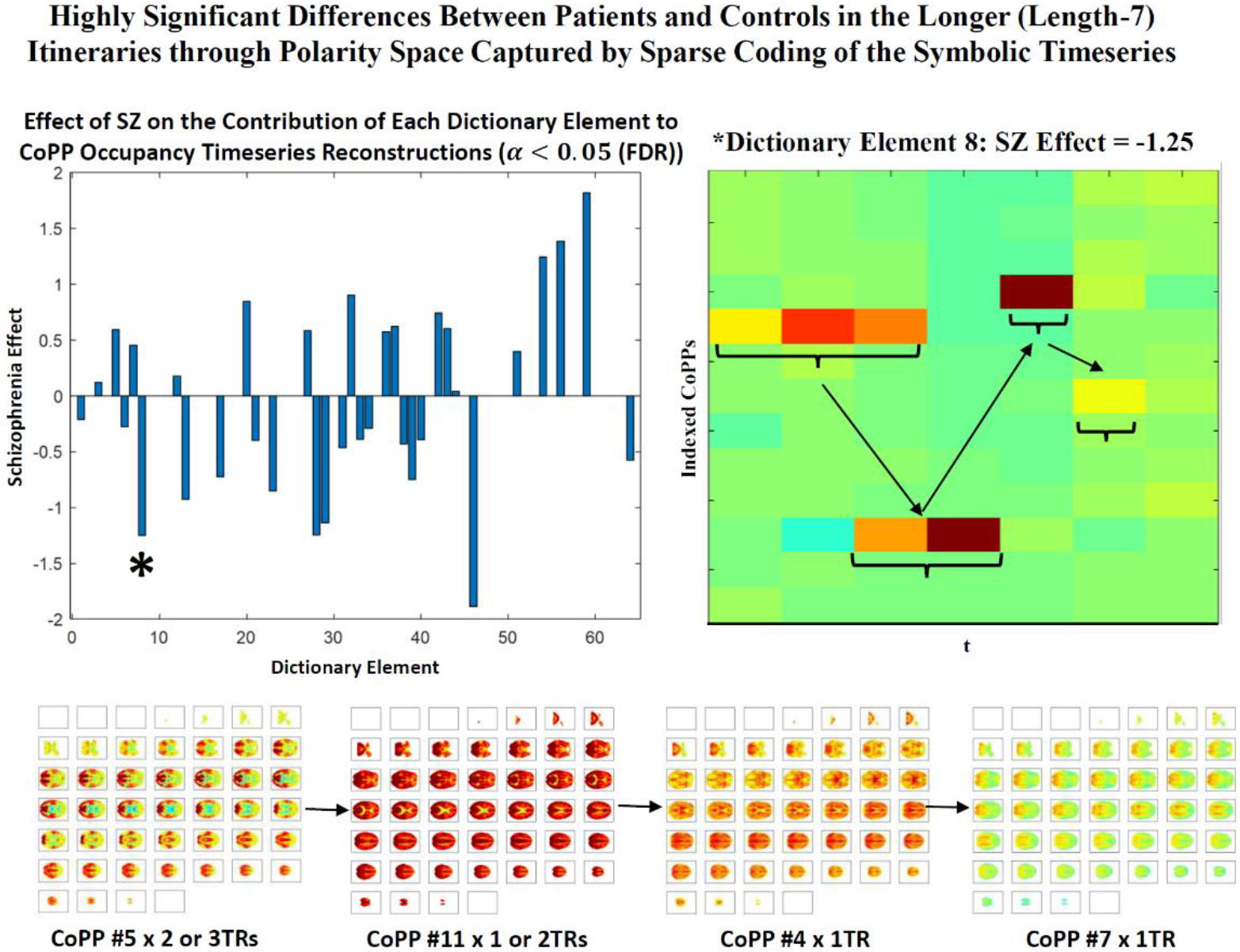
(Top Left) Significant ***p*** < **0.05** (FDR)) effects of schizophrenia on the contributions of each of 64 elements of a13-channel 1-dimensional sparse dictionary of **7**TR temporally convolutional encodings of the original timeseries of CoPP occupancies [17]. Since the CoPP indices are merely arbitrary labels and not numerically meaningful the timeseries are transformed into stacked binary timeseries, each of the univariate timeseries coded as ‘1’ at timepoints when that CoPP is occupied, and ‘0’otherwise. Multichannel sparse temporally convolutional coding represents this timeseries in the form of additive (in this case the coefficients are constrained to be non-negative) contributions of each dictionary element at each timepoint of the input timeseries. The contributions of the different dictionary elements obtained in this way differ very significantly between patients and controls, even after correction for multiple comparisons; (Top Right) One dictionary element (element #8, indicated with a large asterisk in top left panel) that is significantly less present in the representations of patient CoPP occupancy timeseries than the representations of CoPP occupancy timeseries of controls; (Bottom Row) Actual CoPP states in order of occupancy represented as strongly present in the dictionary element displayed in top right panel; this element is showing high probability of being in CoPP #5 (highly structured, highly positively polarized state) for 2 or 3 TRs, followed by a high probability of being next in CoPP #11 (strongly pervasively positively polarized) for 2 or 3TRs, then high probability of next being in CoPP #4 (pervasively positively polarized for short period, then transitioning to CoPP #7 (broadly neutral/non-polarized with some light positive polarization).

## I DISCUSSION

We report results about large-scale spatial BOLD patterns formed by voxel activation timeseries that have each been normalized with respect to their own means and variances (i.e., individually *z-*scored). Our findings indicate that the healthy brain, through some combination of neural, vascular and neurovascular coupling factors, is strongly characterized by the degree to which it exhibits periods of spatially pervasive elevated (resp. depressed) vn-BOLD activation (see Figure 6 and Figure 11). The relative inability of schizophrenia patients to organize this level of pervasive polarization is extremely statistically significant (*p* < 10^−11^). There are also significant differences between healthy and patient populations in the spatial distribution of voxel participation in these states of pervasive brain polarization with stronger structured regions of highly participating voxels among healthy controls (Figure 9). This large-scale brain *polarization* phenomenon exhibits tractable and clinically relevant interactions with familiar connectivity-based targets of fMRI analysis: functional network spatial maps, functional network connectivity and dynamic functional network connectivity. The voxels most drawn into whole-brain polarization states distribute differently over group-level GICA spatial maps in patients and controls (see Figure 10): auditory, visual and sensorimotor networks contain more voxels with high polarity participation rates in controls than patients whereas subcortical and cerebellar networks contain more voxels with high polarity participation rates in patients than controls. There are also significant relationships between polarization and short-timescale functional network connectivity states (Figure 7), relationships that are predominantly unmediated by clinical status with the single exception of the highly modularized state with negative correlations between default-mode and other networks. This dynamic connectivity state is significantly more occupied by controls under conditions of higher than average polarization, while its occupancies by patients are both more numerous and occur under conditions of lower than average levels of whole-brain polarization. In traditional “static” FNC the largely unmediated relationship between polarization and the structure of network correlations/anti-correlations persists (Figure 8). Polarization both guides/constrains and is built from underlying networks and their connectivity. We show that the phenomenon of dynamic, large-scale spatially pervasive polarization in vn-BOLD fMRI is strongly suppressed among those with diagnosed schizophrenia however, with significant intensification of the suppressive effect among patients with higher positive symptomology scores (Figure 6). Positive symptoms include the core psychotic symptoms of schizophrenia, delusions and hallucinations. We have also shown that patients and controls exhibit very different patterns of voxel participation in the polarized TRs (Figure 9). The difference in spatial patterns of “polarity participation” between patients and controls is also present at the network scale. Again it is not clear whether this arises more from voxel effects aggregating into network effects or whether coherent network behavior is shaping most of the voxel participation patterns. Drilling into the transient spatial patterns of discretized voxel IALs yields a set of 13 whole brain co-polarization patterns (CoPPs) that capture, among other things, multiple ways the collection of voxels simultaneously at a given intrinsic activation level (IAL) distribute spatially at timepoints when the whole brain occupies the corresponding dPR state (e.g. voxels with IAL=1 when the whole brain is occupying the polarized-high dPR state). The most strongly polarized CoPPs are significantly more occupied by controls than patients which is consistent with results from the coarse polarity summary provided by dPR state occupancies (Figure 11). However, with CoPPs, we also see group-level evidence of a *healthy* circuit itinerary in polarity space that cycles through the maximally positively polarized CoPP, progressing through four CoPPs with increasing presence and strength of negatively polarized regions until reaching the maximally negatively polarized CoPPs and returning again to the maximally positively polarized CoPP through three intermediate transition states. Using the group-averaged transition probabilities for healthy subjects, i.e. tracing from the most strongly positively polarized CoPP using the highest group-average probability forward-time transition (to a *different state*) and continuing in this manner leads through the most strongly negatively polarized CoPP and then to back to the most strongly positively polarized CoPP again, presenting a group-level most probable *healthy* itinerary through polarity space accounting for 9 of the 13 CoPPs, including the most strongly polarized states. For schizophrenia patients the group-level most probable forward-time trajectory out of the maximally positively polarized CoPP never recovers a significant level of polarity, either positive or negative, instead alternating between two non-polarized states (Figure 12).

There are many neurobiological and vascular factors that could be contributing to the observed tendency of the healthy brain to organize itself toward globally polarized vn-BOLD states, and multiple pathways by which this convergence fails to materialize as fully or as frequently among schizophrenia patients. The inability to “fully polarize” has both locally structured and global manifestations (note that both the strongly polarized structured CoPPs and the strongly pervasively polarized CoPPs in Figure 11 are significantly more occupied by controls, and further, in Figure 12, that strongly polarized structured states are precursors to strongly pervasively polarized state in the attracting orbit for healthy controls). This is simply to point out that the phenomenon we have observed manifests on both local and more global scales, and its disruption in schizophrenia patients is evident at both of these scales. In sum, the proposed approach provides a powerful new perspective on the disruption of brain function, represents a novel way of studying resting fMRI data that does not rely on the standard network-based paradigm, and has the potential to lead to additional biomarkers of brain disease.

References

